# Back to the Future- Unleashing your cytometer’s spectral potential

**DOI:** 10.1101/2022.12.21.521417

**Authors:** Christopher Hall, Hanan Ibrahim, Sam Thompson, Philip S Hobson, Jo-Anne Crofts, Peter Nobes, Steven Lim, Tony Burpee, Rachael V Walker

## Abstract

With the recent growth in spectral flow cytometry many laboratories are investing in new spectral flow cytometers in order to maximise the information gathered about every cell. This study hypothesised that traditional cytometers already within many laboratories may be used as spectral cytometers and have shown using a range of different cytometers that data acquired may be unmixed after acquisition.

## Introduction

In the past couple of years there has been an increase in the number of spectral flow cytometers (also known as Full Spectrum) within laboratories including Shared Resource Laboratories and bio-tech companies, with over 1000 systems being installed [1]. Spectral flow cytometers allow scientists to use more fluorochromes in their experiments, remove auto-fluorescent (AF) signals, and allows live unmixing or subsequent reanalysis at a future date [2, 3]. These benefits are realised through the mathematics used to deconvolute component signals, known as unmixing. The essential characteristic of spectral flow cytometry is the acquisition of data with more parameters than fluorochromes (an overdetermined system). This suggests that a traditional (also referred to as conventional) flow cytometer with all its detectors enabled can use the same deconvolution mathematics to obtain the same or similar results when compared to a spectral system.

Each spectral flow cytometer uses a proprietary method to perform unmixing, so to allow comparison of traditional and spectral flow cytometer data a shared method was developed. As each flow cytometer contained a different number of detectors, together with associated optical filters, the shared method must be able to unmix data from any type of instrument. As a spectral response is predicated on the use of an over-determined system each of the panels used contained fewer fluorochromes than the instrument with the least number of detectors, in this case the Attune with 14 detectors.

The hypothesis of this study aims to show that traditional cytometers can be used with post-acquisition spectral unmixing giving comparable results to spectral instruments.

## Methods

### Sample preparation

Mouse splenocytes were fixed (BD CellFIX, BD Biosciences, 340181) and washed in Flow Cytometry Staining Buffer (eBioscience, 00-4222-26). Cells were preincubated in FC Blocking solution (TruStain FcX™ PLUS, Biolegend, 156603) and Super Bright Staining Buffer (Invitrogen, SB-4401-42) before staining for 45 minutes with following nine fluorochrome panel (all Invitrogen); CD11b FITC (11-0112-41), CD19 eFluor450 (48-0193-82), CD25 APC (17-0251-82), CD3e PerCP-Cy5.5 (45-0031-82), CD4 eFluor506 (69-0042-82), CD45.2 APC-eFluor780 (47-0454-82), CD8a Super Bright 780 (78-0081-82), NK1.1 Super Bright 600 (63-5941-82), and Ly6G AlexaFluor 700 (56-5931-82). Cells were washed twice with Flow Cytometry Staining Buffer and resuspended in 1ml of the same buffer before running on the flow cytometers.

### Data acquisition

Samples were acquired over two days on seven flow cytometry analysers. Two spectral flow cytometers; Sony ID7000 (5 laser and 159 fluorescent detectors (5L/159)) and Cytek Aurora (5L/64), and six traditional instruments; Becton Dickinson (BD) Symphony (5L/30), Beckman Coulter Cytoflex LX (6L/27), Agilent Penteon (5L/30), Bio-Rad ZE5 (5L/30) BD Fortessa (5L/18), and Thermo Fisher Attune CytPix (4L/14). Full instrument configurations are shown in Supp.1 The full spectrum instruments and Penteon were set up using the manufacturer’s recommended voltages (e.g. CytekAssaySettings) and the other traditional instruments were set up by adjusting the detector voltages to place the negative cells at similar median fluorescent intensities (MFI) towards the lower end of the detection range.

### Data Analysis

All data was unmixed in R using the package flowUnmix (github.com/hally166/flowUnmix). flowUnmix which uses the base R function lsfit() to determine the event coefficients based on the single colour controls specified. It has a rudimentary automated population selection method for the controls (taking the top 200 brightest events), but if this fails to detect the spectra, further R code is required to select the positive population (most often with low brightness signals). In some cases, manually gated FCS data was used to choose which cells were selected to form the spectra, the data is then saved as a gated FCS file and used in the flowUnmix R code. Code used for unmixing can be found in the supplementary information (supp figure 1) and on github (github.com/hally166/Unmix_all_data_paper).

Data was additionally analysed and visualised in FlowJo using both AutoSpill [4] and traditional compensation as defined by Bagwell and Adams [5]. All code, packages, and FCS data is freely available from GitHub and AppliedCytometry/FlowRepository

An R package (flowSpectrum) for visualising the spectra of fluorochromes from any instrument has also been designed and was used to check the spectral response from each machine and fluorochrome and is freely available on https://github.com/hally166.

Instrument performance data was additionally analysed, visualised and verified in VenturiOne (www.appliedcytometry.com) (supplementary figure 2).

## Results

With all the detectors enabled and the unstained sample MFI set to a consistent low level (following each instruments manufacturer recommended technique), each instrument produces a unique spectrum for every fluorochrome. These spectra have varying degrees of resolution based on the number of detectors present on the instrument. Figure 1 shows the normalised spectrum of PerCP-Cy5.5 from nine flow cytometers all of which, except the Attune which does not have a UV laser, show the characteristic five emission peaks of PerCP-Cy5.5 thanks to its broad excitation and emission characteristics. The greater the number of detectors the more detailed the emission profile. It was found that even the instrument with the lowest number of detectors (Attune) produced unique spectra for every fluorochrome tried (Supplementary figure 1). This means that as long as overdetermined data is used (data with more detectors than fluorochromes), it is possible to use deconvolution methods, e.g., least squares estimation, to unmix the data (it also possible with the same number of detectors and fluorochromes, but this is considered to be traditional compensation).

**Figure 1:**
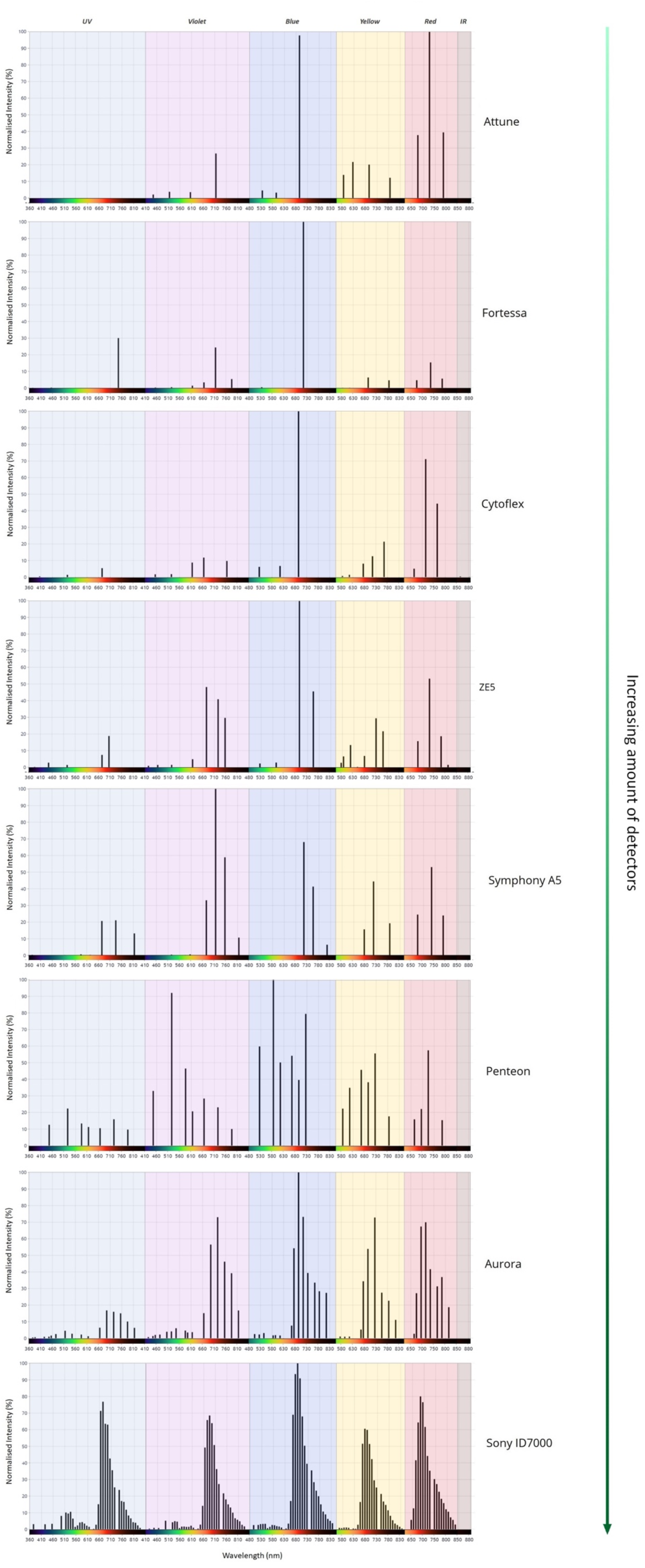
Normalised spectral plots of the fluorochrome PerCP-Cy5.5 on mouse splenocytes produced using nine instruments; two full spectrum (ID7000 and Aurora) and six “traditional” instruments (Symphony, Cytoflex, Penteon, ZE5, Fortessa, and Attune). Spectra are ordered by excitation laser and emission from low to high.

The data from all instruments was unmixed using the R package flowUnmix to allow more consistent comparison, as manufacturer unmixing in the acquisition software may introduce other factors to improve visualisation and resolution, such as weightings and transformations. The coefficients which comprise the unmix matrix used by the R package were independently validated using VenturiOne (Supplementary figure 2). The manufacturer unmixed data is included in the data repository. Figure 2 shows the unmixed data from the seven instruments visualised in FlowJo (BD). There was a high degree of uniformity in the population distributions across all the instruments, except the Cytoflex and Attune, both of which showed poor separation of the CD4 eFluor506 and CD8 Super Bright 780 signals, however the percentages were still consistent with the other instruments. It should be noted that the Attune does not have a detector to optimally observe the Super Bright 780 signal however, this fluorochrome was resolved through unmixing (see Attune CD8 data as indicated in red square in Fig. 2).

**Figure 2:**
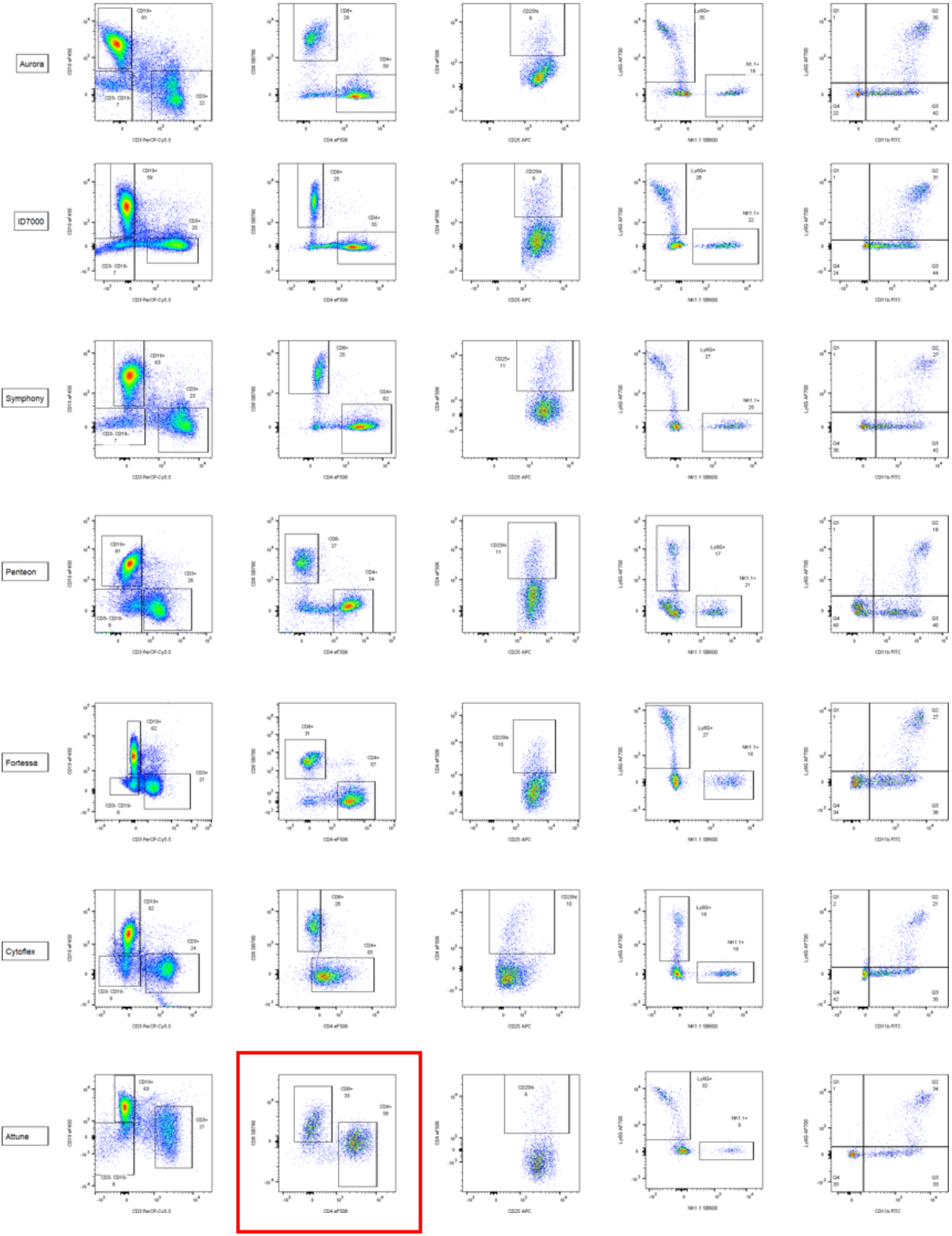
A nine parameter experiment was performed on seven instruments, two full spectrum and five traditional. Each was unmixed in R using the package flowUnmix and visualised in FlowJo. Each row is one instrument and each column is one bivariant plot described on the top row. Gates were drawn based on the unstained control and axis labels are generated from the control file names. All samples were gated on scatter and CD42.2. Red box around Attune CD8 vs CD4 plot shows the Superbright 780 can be observed despite there not being a traditional filter for this fluorochrome.

To test if least squares unmixing performs as well as traditional compensation we calculated the staining index of the fluorochromes using unmixing, AutoSpill and traditional compensation. Table 1 shows the staining index of the unmixed data and the positive and negative fold change when using Autospill and compensation. Signal separation by staining index was usually less with unmixing, except with the NK1.1 Super Bright 600 which showed better separation. It is difficult to make any broad judgements as performance was dependent on the instrument and the fluorochrome being used.

**Table 1:**
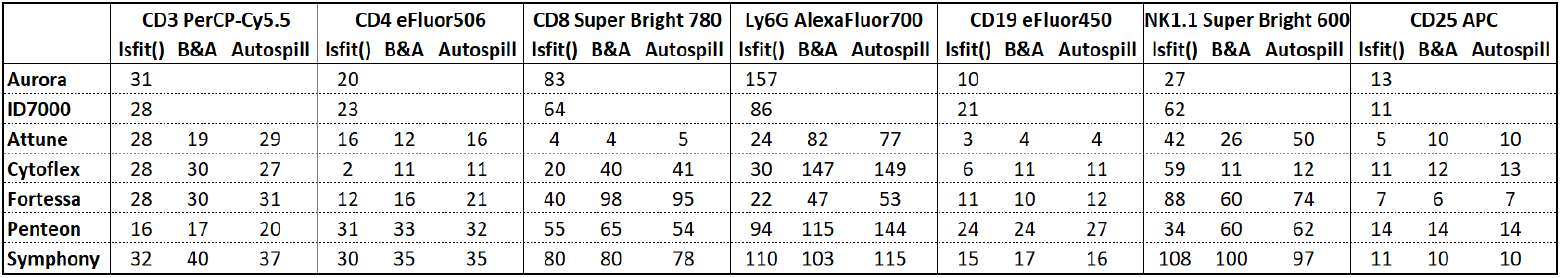
The staining index of the data when using unmixing (lsfit()), traditional compensation (as derived by Bagwell & Adams(B&A)), and Autospill

## Discussion

The “sold as” full spectrum flow cytometer segregates the spectrum into differently sized segments, the same method is of course used in a traditional instrument. This led to the hypothesis that all flow cytometers can be treated as a full spectral machine, because it is the unmixing mathematics that generates the benefits of the technology. This paper shows that data from all the instruments that we tested can be successfully unmixed and the data resolution can be comparable with spectral machines. The R based tools, flowUnmix and flowSpectrum, have been made available to allow others to unmix their own data.

There are several ways to set up the instrument to attain a clean fluorochrome spectra for unmixing. Both the Aurora and the ID7000 use an optimised base setting as a starting point that does not have a fixed negative fluorescence, but an optimised one based on the most common fluorochromes and known instrument performance. Over multiple unmixings on the instruments the most common adjustment involved PerCP-Cy5.5 and BV711. Both of which often required minor alterations in detector settings to prevent them from distorting the unmixing by being overrepresented in the incorrect channel and reducing the resolution of other fluorochromes. It is expected that optimised settings, similar to those on the ID7000 and Aurora, would result in better population resolution, but this is yet to be confirmed.

There are differences in the qualitative nature of the data from each instrument. This can be attributed to the number and type of detectors being employed. It has been suggested that the more overdetermined the system, the better the deconvolution. This study cannot ascertain if there is an ideal level of overdetermination or filter setup because there are too many confounding factors, most notably, the choice of fluorochrome. However, it is clear from the data that any overdetermined system can be unmixed, whether it is “sold as” full spectral or not. It should be noted that the Aurora and ID7000 instruments, used as the ground truth controls here, produce data with differing appearance, and to a small degree, differing population proportions. This has been known for a long time[6] and is the reason normalisation algorithms exist to allow the comparison of data across instruments. The nature of unmixing will add to this complexity as there is no “correct” answer just the best fit case for each deconvolution coefficient. The coefficients can be adjusted depending on the fitting method used. Sony have published the fact that they use a weighting in their unmixing to better fit the data produced by their instrument.[7]

In this comparison with traditional compensation and Autospill, an interesting side effect of Autospill that was not discussed in the original paper by Rocca et al was discovered. That is (in the FlowJo implementation) that the incremental adjustment step that corrects for compensation issues after the first linear compensation can produce large negative values in the compensation matrix, effectively adding signal to the detectors that was not actually there. This was evident on the samples, such as the ZE5 example, where the experiment was not set up optimally for traditional compensation (I.e., fluorochrome brightest in the correct detector). This could possibly be seen as analogous to the least squares fitting that is done during unmixing where the regression chooses the best fit of the data at a point irrespective of the whether it is only removing data, as would be the case in traditional compensation.

A surprising observation that the data revealed was the ability of the Attune to resolve the Superbright 780 fluorophore, considering this machine did not have a dedicated filter setup to detect this fluorochrome. This introduces a range of possibilities for fluorochrome combinations in panel design that were not previously available on traditional machines.

By unmixing all the instruments using the same least squares regression it is possible to objectively compare the performance of each instrument. By comparing the flowUnmix unmixed data with the manufacturer’s unmixed data, there appear to be differences (supp data). This suggests that there may be a possibility of adjusting the unmixing to better fit the raw data. For example, unmixing based on the fluorophores being used or the scientific question being asked. This has been common practice in microscopy for many years [8, 9] and there are opportunities to do the same in flow cytometry. There is also the possibility to extend the benefits discussed of spectral flow cytometry, such as autofluorescence subtraction, to traditional instruments.

## Conclusions

This study has shown that traditional cytometers can be used as spectral flow cytometers. This allows for each fluorochrome to be measured across all lasers and complete spectrum of each dye to be collected and unmixed. Resulting in the ability to use fluorophores that could not be previously identified on that flow cytometer.

## Acknowledgements

We would like to thank Dr Bethan Jones, (Sr Product Owner Thermo-Fisher, Eugene) for the donation of the antibodies used in this study. Also, thanks to Agilent for giving us access to the Penteon. Thanks to Babraham Institute and the Francis Crick Institute Flow Cytometry Facilities for supporting and facilitating this work. The Babraham Institute Flow Cytometry facility was supported by the Babraham Institute’s UKRI-BBSRC Core Capability Grant.

