## Supplementary files (fig1-3) for "Back to the Future- Unleashing your cytometer’s spectral potential"

**Supplementary Information –**

Supplementary Figure 1: Example code for unmixing the Attune data:

```
# R 4.1.1 (2022)
# Install the required packages.  You may also need to install devtools,
biocManager,
# flowCore, ggcyto, etc. Pay attention to the warning messages

devtools::install_github("hally166/flowUnmix")
biocManager::install("ggcyto")
library(flowUnmix)
library(ggcyto)

# Attune (Babraham) -- mouse (no SB780 detector)

files<-list.files("C:/Attune 07OCT22/Mouse", pattern = "fcs",
full.names=TRUE)
files #check files
controlfilesFS<-read.flowSet(files[c(1:7,9:10)]) #select the controls
unstainedcontrol<-read.FCS(files[11]) #specify the unstained
filetounmix<-read.flowSet(files[8]) #specify the file(s) to unmix

# Fails to correctly select APC signature - gate manually

samp <- controlfilesFS[[3]]
colnames(samp)
p <- autoplot(samp, "RL1-A")
p + scale_x_logicle()
p + scale_x_logicle() + geom_vline(xintercept = c(10000,69000))
rectGate <- rectangleGate(filterId="apc","RL1-A"=c(10000,69000))
controlfilesFS[[3]]<-Subset(samp, rectGate)

# Use flowUnmix to unmix the data

flowUnmix(fs=filetounmix, cs=controlfilesFS, unstained =unstainedcontrol,
guessPop = TRUE, popCheck = TRUE)
```

Supplementary Figure 2: Graphics showing data unmixed using R method and then validated using VenturiOne software.

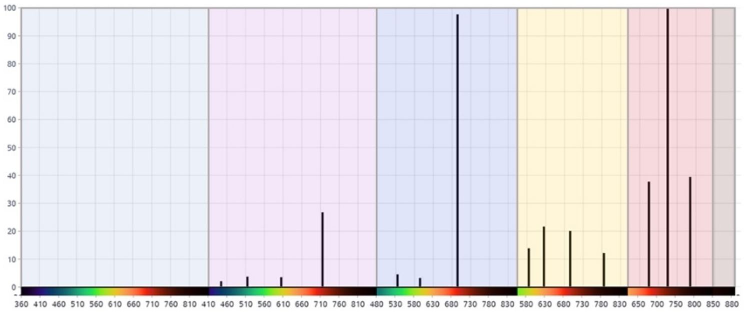

Attune

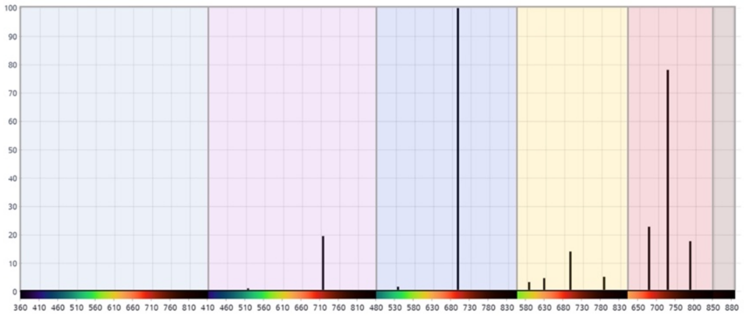

VenturiOne Attune Data

Supplementary Figure 3: Instrument details for cytometers used in this study

| **Laser (nm)** | **Power (mW)** |
|---|---|
| 355 | 24 |
| 405 | 50 |
| 488 | 50 |
| 561 | 50 |
| 640 | 40 |

| **Laser (nm)** | **Bandpass Filter** |
|---|---|
| 355 | 379/28 |
| | 450/50 |
| | 530/30* |
| | 670/25* |
| | 740/35 |
| | 820/60* |
| 405 | 450/50 |
| | 525/50 |
| | 610/20 |
| | 660/20 |
| | 710/50 |
| | 780/60 |
| 488 | 530/30 |
| | 710/50 |
| 561 | 585/15 |

| | |
|---|---|
| | 610/20 |
| | 695/40 |
| | 780/60 |
| 640 | 670/14 |
| | 730/45 |
| | 780/60 |

Figure 3.1. BD LSRFortessa™ Cell Analyzer equipped lasers and power (L) and optical configuration (R).

| Laser (nm) | Power (mW) |
|---|---|
| 355 | 50 |
| 405 | 100 |
| 488 | 50 |
| 561 | 50 |
| 640 | 60 |

| Laser (nm) | Bandpass Filter |
|---|---|
| 355 | 387/11 |
| | 447/60 |
| | 525/50 |
| | 670/30 |
| | 700LP |
| 405 | 420/10 |
| | 460/22 |
| | 525/50 |
| | 615/24 |
| | 670/30 |
| | 720/60 |
| | 750LP |
| 488 | 488/10 |
| | 525/35 |
| | 593/52 |
| | 692/80 |
| | 750LP |
| 561 | 577/15 |
| | 589/15 |
| | 615/24 |
| | 640/20 |
| | 670/30 |
| | 720/60 |
| | 750LP |
| 640 | 670/14 |
| | 720/60 |
| | 775/50 |
| | 800LP |

Figure 3.2. BioRad ZE5 (Yeti) Cell Analyzer equipped lasers and power (L) and optical configuration (R).

| Laser (nm) | Power (mW) |
|---|---|
| 405 | 100 |
| 488 | 100 |
| 561 | 100 |
| 642 | 140 |

| Laser (nm) | Bandpass Filter |
|---|---|
| 405 | 440/50 |
| | 512/25 |
| | 603/48 |
| | 710/50 |
| 488 | 488/10+0D2 |
| | 530/30 |
| | 590/40 |
| | 695/40 |
| 561 | 585/16 |
| | 620/15 |
| | 695/40 |
| | 780/60 |
| 640 | 670/14 |
| | 720/30 |
| | 780/60 |

Figure 3.3. Thermo Fisher Attune™ CytPix™ Flow Cytometer equipped lasers and power (L) and optical configuration (R)

| Laser (nm) | Power (mW) |
|---|---|
| 349 | 20 |
| 405 | 100 |
| 488 | 100 |
| 561 | 100 |
| 637 | 100 |

| Laser (nm) | Bandpass Filter |
|---|---|
| 349 | 445/45 |
| | 525/45 |
| | 586/20 |
| | 615/20 |
| | 667/30 |
| | 725/40 |
| | 780/60 |
| 405 | 445/45 |
| | 525/45 |
| | 586/20 |
| | 615/20 |
| | 667/30 |
| | 725/40 |
| | 780/60 |
| 488 | 525/45 |
| | 586/20 |
| | 615/20 |
| | 667/30 |
| | 695/40 |

| | |
|---|---|
| | 725/40 |
| 561 | 586/20 |
| | 615/20 |
| | 667/30 |
| | 695/40 |
| | 725/40 |
| | 780/60 |
| 640 | 667/30 |
| | 695/40 |
| | 725/40 |
| | 780/60 |

Figure 3.4. Agilent NovoCyte® Penteon™ Flow Cytometer equipped lasers and power (L) and optical configuration (R).

| Laser (nm) | Power (mW) |
|---|---|
| 355 | 20 |
| 405 | 100 |
| 488 | 50 |
| 561 | 50 |
| 640 | 80 |

| Laser (nm) | Channel | Bandpass Filter |
|---|---|---|
| 355 | UV1 | 372/15 (355) |
| | UV2 | 387/15 (355) |
| | UV3 | 427/15 (355) |
| | UV4 | 443/15 (355) |
| | UV5 | 458/15 (355) |
| | UV6 | 473/15 (355) |
| | UV7 | 514/28 (355) |
| | UV8 | 542/28 (355) |
| | UV9 | 581/31 (355) |
| | UV10 | 612/31 (355) |
| | UV11 | 664/27 (355) |
| | UV12 | 691/28 (355) |
| | UV13 | 720/29 (355) |
| | UV14 | 750/30 (355) |
| | UV15 | 780/30 (355) |
| | UV16 | 812/34 (355) |
| 405 | V1 | 428/15 (405) |
| | V2 | 443/15 (405) |
| | V3 | 458/15 (405) |
| | V4 | 473/15 (405) |
| | V5 | 508/20 (405) |
| | V6 | 525/17 (405) |
| | V7 | 542/17 (405) |
| | V8 | 581/19 (405) |
| | V9 | 598/20 (405) |

| | | |
|---|---|---|
| | V10 | 615/20 (405) |
| | V11 | 664/27 (405) |
| | V12 | 692/28 (405) |
| | V13 | 720/29 (405) |
| | V14 | 750/30 (405) |
| | V15 | 780/30 (405) |
| | V16 | 812/34 (405) |
| 488 | B1 | 508/20 (488) |
| | B2 | 525/17 (488) |
| | B3 | 542/17 (488) |
| | B4 | 581/19 (488) |
| | B5 | 598/20 (488) |
| | B6 | 615/20 (488) |
| | B7 | 660/17 (488) |
| | B8 | 678/18 (488) |
| | B9 | 697/19 (488) |
| | B10 | 717/20 (488) |
| | B11 | 738/21 (488) |
| | B12 | 760/23 (488) |
| | B13 | 783/23 (488) |
| | B14 | 812/34 (488) |
| 561 | YG1 | 577/20 (561) |
| | YG2 | 598/20 (561) |
| | YG3 | 615/20 (561) |
| | YG4 | 660/17 (561) |
| | YG5 | 678/18 (561) |
| | YG6 | 697/19 (561) |
| | YG7 | 720/29 (561) |
| | YG8 | 750/30 (561) |
| | YG9 | 780/30 (561) |
| | YG10 | 812/34 (561) |
| 640 | R1 | 660/17 (640) |
| | R2 | 678/18 (640) |
| | R3 | 697/19 (640) |
| | R4 | 717/20 (640) |
| | R5 | 738/21 (640) |
| | R6 | 760/23 (640) |
| | R7 | 783/23 (640) |
| | R8 | 812/34 (640) |

Figure 3.5. Cytek® Aurora equipped lasers and power (L) and optical configuration (R)

| Laser (nm) | Power (mW) |
|---|---|
| 355 | 60 |
| 405 | 100 |
| 488 | 100 |
| 561 | 150 |
| 633 | 100 |

| Laser (nm) | Bandpass Filter |
|---|---|
| 355 | 379/28 |
| | 515/30 |
| | 580/20 |
| | 605/20 |
| | 670/25 |
| | 735/30 |
| | 810/40 |
| 405 | 431/28 |
| | 525/50 |
| | 586/15 |
| | 605/40 |
| | 677/20 |
| | 710/20 |
| | 750/30 |
| | 780/60 |
| 488 | 488/10 |
| | 530/30 |
| | 610/20 |
| | 670/30 |
| | 710/50 |
| | 780/60 |
| 561 | 586/15 |
| | 610/20 |
| | 670/30 |
| | 710/50 |
| | 780/60 |
| 633 | 670/30 |
| | 730/45 |
| | 780/60 |

Figure 3.6 BD FACSymphony™ A5 Cell Analyzer equipped lasers and power (L) and optical configuration (R).

| Laser (nm) | Power (mW) |
|---|---|
| 355 | 50 |
| 405 | 100 |
| 488 | 150 |
| 561 | 100 |
| 637 | 140 |

| Laser (nm) | Label | Bandpass Filter |
|---|---|---|
| 355 | CH1 | 350.5-390 |
| | CH2 | 418.5-446 |
| | CH3 | 446-472.5 |
| | CH4 | 493.9-506.1 |
| | CH5 | 506.1-518.3 |

| | | |
|---|---|---|
| | CH6 | 518.3-530.3 |
| | CH7 | 530.3-542.3 |
| | CH8 | 542.3-554.2 |
| | CH9 | 554.2-566.1 |
| | CH10 | 566.1-577.8 |
| | CH11 | 577.8-589.5 |
| | CH12 | 589.5-601.1 |
| | CH13 | 601.1-612.6 |
| | CH14 | 612.6-624 |
| | CH15 | 624-635.3 |
| | CH16 | 635.3-646.6 |
| | CH17 | 646.6-657.7 |
| | CH18 | 657.7-668.8 |
| | CH19 | 668.8-679.8 |
| | CH20 | 679.8-690.8 |
| | CH21 | 690.8-701.6 |
| | CH22 | 701.6-712.4 |
| | CH23 | 712.4-723 |
| | CH24 | 723-733.6 |
| | CH25 | 733.6-744.2 |
| | CH26 | 744.2-754.6 |
| | CH27 | 754.6-765 |
| | CH28 | 765-775.2 |
| | CH29 | 775.2-785.4 |
| | CH30 | 785.4-795.5 |
| | CH31 | 795.5-805.5 |
| | CH32 | 805.5-815.5 |
| | CH33 | 815.5-825.3 |
| | CH34 | 825.3-835.1 |
| | CH35 | 835.1-844.7 |
| 405nm | CH1 | 413.6-438 |
| | CH2 | 438-459 |
| | CH3 | 459-480.2 |
| | CH4 | 493.9-506.1 |
| | CH5 | 506.1-518.3 |
| | CH6 | 518.3-530.3 |
| | CH7 | 530.3-542.3 |
| | CH8 | 542.3-554.2 |
| | CH9 | 554.2-566.1 |
| | CH10 | 566.1-577.8 |
| | CH11 | 577.8-589.5 |
| | CH12 | 589.5-601.1 |

| | | |
|---|---|---|
| | CH13 | 601.1-612.6 |
| | CH14 | 612.6-624 |
| | CH15 | 624-635.3 |
| | CH16 | 635.3-646.6 |
| | CH17 | 646.6-657.7 |
| | CH18 | 657.7-668.8 |
| | CH19 | 668.8-679.8 |
| | CH20 | 679.8-690.8 |
| | CH21 | 690.8-701.6 |
| | CH22 | 701.6-712.4 |
| | CH23 | 712.4-723 |
| | CH24 | 723-733.6 |
| | CH25 | 733.6-744.2 |
| | CH26 | 744.2-754.6 |
| | CH27 | 754.6-765 |
| | CH28 | 765-775.2 |
| | CH29 | 775.2-785.4 |
| | CH30 | 785.4-795.5 |
| | CH31 | 795.5-805.5 |
| | CH32 | 805.5-815.5 |
| | CH33 | 815.5-825.3 |
| | CH34 | 825.3-835.1 |
| | CH35 | 835.1-844.7 |
| 488nm | CH4 | 493.9-506.1 |
| | CH5 | 506.1-518.3 |
| | CH6 | 518.3-530.3 |
| | CH7 | 530.3-542.3 |
| | CH8 | 542.3-554.2 |
| | CH9 | 554.2-566.1 |
| | CH10 | 566.1-577.8 |
| | CH11 | 577.8-589.5 |
| | CH12 | 589.5-601.1 |
| | CH13 | 601.1-612.6 |
| | CH14 | 612.6-624 |
| | CH15 | 624-635.3 |
| | CH16 | 635.3-646.6 |
| | CH17 | 646.6-657.7 |
| | CH18 | 657.7-668.8 |
| | CH19 | 668.8-679.8 |
| | CH20 | 679.8-690.8 |
| | CH21 | 690.8-701.6 |
| | CH22 | 701.6-712.4 |

| | | |
|---|---|---|
| | CH23 | 712.4-723 |
| | CH24 | 723-733.6 |
| | CH25 | 733.6-744.2 |
| | CH26 | 744.2-754.6 |
| | CH27 | 754.6-765 |
| | CH28 | 765-775.2 |
| | CH29 | 775.2-785.4 |
| | CH30 | 785.4-795.5 |
| | CH31 | 795.5-805.5 |
| | CH32 | 805.5-815.5 |
| | CH33 | 815.5-825.3 |
| | CH34 | 825.3-835.1 |
| | CH35 | 835.1-844.7 |
| 561nm | CH10 | 566.1-577.8 |
| | CH11 | 577.8-589.5 |
| | CH12 | 589.5-601.1 |
| | CH13 | 601.1-612.6 |
| | CH14 | 612.6-624 |
| | CH15 | 624-635.3 |
| | CH16 | 635.3-646.6 |
| | CH17 | 646.6-657.7 |
| | CH18 | 657.7-668.8 |
| | CH19 | 668.8-679.8 |
| | CH20 | 679.8-690.8 |
| | CH21 | 690.8-701.6 |
| | CH22 | 701.6-712.4 |
| | CH23 | 712.4-723 |
| | CH24 | 723-733.6 |
| | CH25 | 733.6-744.2 |
| | CH26 | 744.2-754.6 |
| | CH27 | 754.6-765 |
| | CH28 | 765-775.2 |
| | CH29 | 775.2-785.4 |
| | CH30 | 785.4-795.5 |
| | CH31 | 795.5-805.5 |
| | CH32 | 805.5-815.5 |
| | CH33 | 815.5-825.3 |
| | CH34 | 825.3-835.1 |
| | CH35 | 835.1-844.7 |
| 637nm | CH17 | 646.6-657.7 |
| | CH18 | 657.7-668.8 |
| | CH19 | 668.8-679.8 |

| | |
|---|---|
| CH20 | 679.8-690.8 |
| CH21 | 690.8-701.6 |
| CH22 | 701.6-712.4 |
| CH23 | 712.4-723 |
| CH24 | 723-733.6 |
| CH25 | 733.6-744.2 |
| CH26 | 744.2-754.6 |
| CH27 | 754.6-765 |
| CH28 | 765-775.2 |
| CH29 | 775.2-785.4 |
| CH30 | 785.4-795.5 |
| CH31 | 795.5-805.5 |
| CH32 | 805.5-815.5 |
| CH33 | 815.5-825.3 |
| CH34 | 825.3-835.1 |
| CH35 | 835.1-844.7 |

Figure 3.7. Sony ID7000™ Spectral Cell Analyzer equipped lasers and power (L) and optical configuration (R).

| Laser (nm) | Power (mW) |
|---|---|
| 355 | 20 |
| 405 | 80 |
| 488 | 50 |
| 561 | 30 |
| 633 | 50 |
| 805 | 60 |

| Laser (nm) | Label | Bandpass filter |
|---|---|---|
| 355 | UV405 | 405/30 |
| | UV525 | 525/40 |
| | UV675 | 675/30 |
| 405 | Scatter | 405/10 |
| | V450 | 450/45 |
| | V525 | 525/40 |
| | V610 | 610/20 |
| | V660 | 660/10 |
| | V763 | 763/43 |
| 488 | SSC | 488/8 |
| | B525 | 525/40 |
| | B610 | 610/20 |
| | B690 | 690/50 |
| 561 | | 561/6 |
| | Y585 | 585/42 |
| | Y610 | 610/20 |
| | Y675 | 675/30 |
| | Y710 | 710/50 |
| | Y763 | 763/43 |

| | | |
|---|---|---|
| 633 | | 638/6 |
| | R660 | 660/10 |
| | R712 | 712/25 |
| | R763 | 763/43 |
| 805 | IR840 | 840/20 |
| | IR885 | 885/40 |

Figure 3.8. Beckman Coulter CytoFLEX Flow Cytometer equipped lasers and power (L) and optical configuration (R).
